# Epigenome-Based Drug Repositioning in Acute Myeloid Leukemia

**DOI:** 10.1101/152157

**Authors:** Adam S Brown, Chirag J Patel

## Abstract

**Background:** Repositioning approved drugs for the treatment of new indications is a promising avenue to reduce the burden of drug development. Most currently available computational methods based on molecular evidence can only utilize gene expression for repositioning despite a growing interest in the epigenome in human disease. We recently described a novel repositioning method, ksRepo, that enables investigators to move beyond microarray-based gene expression and utilize a variety of other sources of molecular evidence, such as DNA methylation differences.

**Methods:** We downloaded differential DNA methylation data from two publicly available acute myeloid leukemia (AML) datasets, a cancer with known, extensive epigenomic perturbations. We consolidated CpGs-level to non-directional gene-level differential methylation using Brown’s correction to Fisher’s method. We then used ksRepo, which ignores directionality in disease- and gene-drug associations, to mine the resulting prioritized gene lists and and the Comparative Toxicogenomics Database (CTD) for predicted repositioning candidates.

**Results:** We successfully recovered four compounds that were significant (FDR < 0.05) in two AML datasets: cytarabine, alitretinoin, panobinostat, and progesterone. Cytarabine is the most commonly used frontline therapy for AML and alitretinoin, panobinostat, and progesterone have all been investigated for the treatment of AML.

**Conclusions.** Combining a method for consolidating CpG methylation to the gene level with ksRepo provides a pipeline for deriving drug repositioning hypotheses from differential DNA methylation. We claim that our platform can be extended to other diseases with epigenetic perturbations and to other epigenomic modalities, such as ChIP-seq.

## INTRODUCTION

The process of nominating new indications for previously approved drugs, known as drug repositioning, has become increasing attractive to industry and academia due to the substantial decrease in risk of unforeseen adverse events associated with compounds with known safety profiles [1–3]. A large number of computational approaches have been developed over the past 10 years that leverage molecular evidence to identify repositioning candidates, such as using differential gene expression [4–7]. Unfortunately, these methodologies are hindered by their need for specific data types, assay platforms, and data formats, preventing investigators from utilizing newer profiling technologies and expanding beyond gene expression. In particular, epigenomics is a key modality that has yet to be broadly utilized, in large part due to the lack of clear directionality at the gene level associated with differential DNA methylation and histone modifications; for instance, intra-exon DNA methylation can lead to upregulation or downregulation of gene expression depending on a variety of factors, including histone modifications [8,9].

Despite this complexity, epigenomic studies have been proposed as a promising data source for driving personalized medicine [10,11], but to our knowledge no methods have been published that allow the direct translation of epigenomic findings to drug repositioning. To address this unmet need, we developed a novel pipeline for identifying candidate repositioning candidates from epigenomic data, and in particular from DNA methylation data. We pair a widely used method to condense CpG-level DNA methylation to gene-level information with our previously described repositioning tool, ksRepo [12]. We demonstrate the utility of our method in AML using two independent genome-wide methylation datasets (see Figure 1). AML is the ideal case-study for our methodology, as recent work has shown that the AML epigenome evolves independently of mutational burden, and, furthermore, patients with AML respond well to DNA demethylating agents (also known as “epigenetic drugs”) [13–16]. Using our pipeline, we recover significance for the most commonly used AML medication, cytarabine, as well as three other candidates with ongoing preclinical research in the AML space.

**Fig 1.**
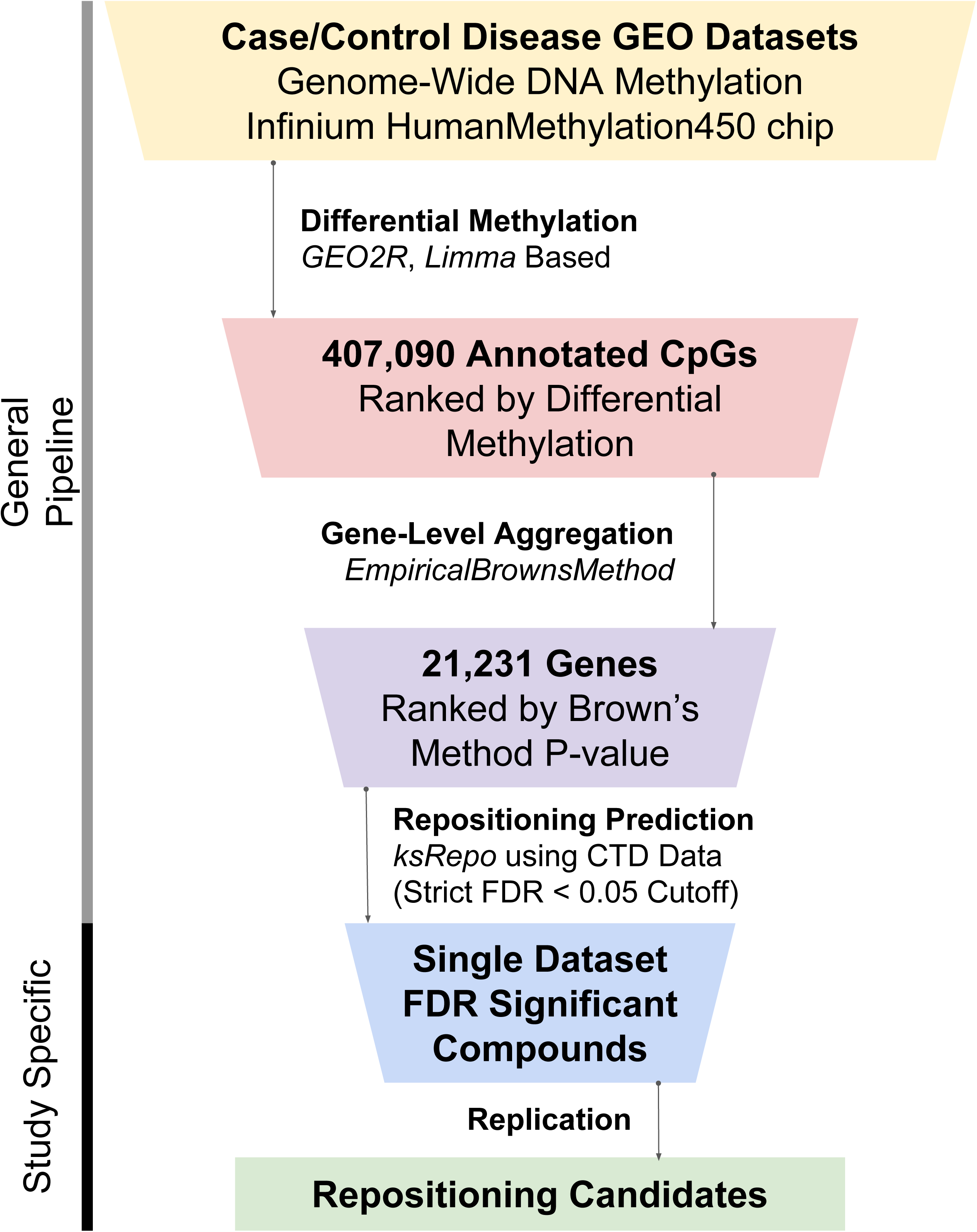
Overview of the general pipeline design and exploratory AML analyses. Our general, DNA methylation-driven repositioning pipeline consists of (1) GEO2R differential methylation analysis, (2) consolidation of CpG-level significance to gene-level significance using Brown’s Method, and (3) ksRepo computational drug repositioning analysis. For this study, we used two publicly available GEO (GSE58477 and GSE63409) datasets and identified four replicated repositioning candidates.

## METHODS

### GEO Dataset Processing

The datasets used in this study are GSE58477 and GSE63409, both of which are Illumina Infinium HumanMethylation450 chip-based (hereafter 450K) studies of AML [17,18]. GSE58477 contained data from a total of 62 patients with cytogenically normal AML and 10 CD34+ normal bone marrow aspirate samples [17]. GSE63409 contained data from a total of 15 AML patients of varying FAB subtype classifications (M0: 1, M1: 2, M2: 2, M5: 5, Not Determined: 5) and five sorted, normal bone marrow aspirate samples [18]. Both GEO datasets were accessed through the NCBI GEO portal and analyzed using the GEO2R tool [19]. GEO2R uses the limma package in R [20], an analytic method originally designed for gene expression, which has been used to detect differential methylation [21,22]. As input for GEO2R, we classified each sample within a GEO series as either normal CD34+ bone marrow aspirate or leukemic blast (as indicated by either lack of engraftment only or by a combination of CD34- and lack of engraftment for GSE58477 and GSE63409 respectively). After classification we were left with 62:10 and 11:5 case:control ratios for GSE58477 and GSE63409 respectively.

### Gene-level differential methylation using Fisher’s Method

To perform methylation-based repositioning, ksRepo requires a list of genes ranked according to their differential methylation. We therefore condensed the GEO2R output from CpG-level to gene-level differential methylation. Consolidating CpG-level significance values to gene-level values requires three steps: (1) identifying CpGs that are annotated to a given gene (obtained from the manufacturer’s website), (2) weighted combining of p-values from CpGs annotated to that gene, and (3) adjusting the resulting, combined p-value to take into account the high degree of correlation between CpGs within a single gene. We performed the weighted combination of p-values, and adjusted them using the *EmpircalBrownsMethod* package in R [23,24]. *EmpiricalBrownsMethod* first combines CpG-level p-values into an omnibus statistic using Fisher’s Method and then calculates an empirical p-value using the correlation structure of the CpGs in each GEO dataset [25]. Using this methodology, we condensed 407,090 individual CpGs into a list of 21,231 genes ranked by their degree of differential methylation (see Figure 1). Some CpGs lacked differential methylation estimates due to poor probe quality; for these CpGs, significance was set as the patient-level median significance for the other CpGs annotated to the same gene. For purposes of differential methylation calculations, all p-values were adjusted using the False Discovery Rate method [26]. After estimating gene-level significance, differentially methylated genes were taken to be those with FDR significance below 0.05. These differentially expressed genes were subjected to pathway enrichment analysis using the PANTHER classification system (PANTHER pathways were used, see [27]). In short, PANTHER contains manually curated pathway annotations for human genes, as well as built-in analytical tools for reporting statistical enrichment using fisher’s exact test and gene-set enrichment; PANTHER was chosen due to the frequency of updates to its pathway annotations. In addition, PANTHER reports all enriched pathways using a strict family-wise error rate correction (to account for spurious findings or type 1 error), with significance deemed as Bonferroni-corrected p-value below 0.05 [28].

### ksRepo repositioning and replication

To determine whether differential methylation-based repositioning is robust, we analyzed each dataset separately using ksRepo (package available for download at https://github.com/adam-sam-brown/ksRepo, and described in [12]). ksRepo uses a modified Kolmogorov-Smirnov (KS) statistic that does not require directionality of effect, and calculates significance using resampling of compound-gene interaction lists. As our database of compound-gene interactions, we used the ksRepo Comparative Toxicogenomics Database (CTD) dataset, which is built into the ksRepo package. The CTD provides a curated resource that links small chemical entities to genes (e.g., gene or protein expression influences) from the scientific literature numerous model organisms and humans [29]. ksRepo contains a subset of the CTD containing human-derived interactions between 1,268 unique drugs and 18,041 unique human genes. Drugs in the ksRepo subset were chosen based on case-insensitive matches between CTD names and names/synonyms for FDA approved drugs downloaded from DrugBank [30]. We combined the ranked gene list from above with the CTD using ksRepo. ksRepo’s output provides both the resampled and false discovery rate (FDR) adjusted p-value [26]. On the study-level, significant compounds were those for which the FDR was less than 0.05. Replicated hits were considered to be those compounds with significance in both methylation studies.

## RESULTS AND DISCUSSION

### Development of a pipeline for epigenome-based repositioning

In this study, we built on our previously developed tool, ksRepo, a drug repositioning methodology that enables investigators to use directionally agnostic, prioritized gene lists to nominate compounds as repositioning candidates. Removing the requirement for directionality of effect or association (e.g. up- or down-regulation of a gene) allows investigators to use complex molecular data, including DNA methylation. In a DNA methylation experiment, a single gene can have several CpG hyper- and hypomethylation events, which would yield both negative and positive associations of DNA methylation with disease. Therefore, it is a challenge to summarize the overall “effect” direction of a gene with both negative and positive associations. As input for ksRepo, we consolidated over 400,000 CpGs to gene-level significance using Fisher’s omnibus statistic and corrected for the high degree of inter-CpG correlation using Brown’s Method (see Figure 1) [25]. Although we describe here a specific use-case of our pipeline for DNA methylation, our methodology can be adapted to other epigenomic signals, such as genome-wide ChIP-seq data from a variety of histone marks and other factors.

### Epigenome-based repositioning using publicly available AML DNA methylation data

To assess the promise of our method, we used two, publicly available 450K DNA methylation datasets from GEO, both in AML [17,18]. Both studies noted clear patterns of hypo- and hypermethylation, with complex alterations in methylation taking place within genes, and especially within Wnt signaling genes and homeobox domain containing genes (HOX genes). Using our gene-level differential methylation consolidation technique, we found that GSE58477 and GSE63409 had 820 and 442 differentially expressed genes respectively (FDR < 0.05). Mirroring the results of the two GEO studies, we found that Wnt signaling genes, were significantly enriched in our method’s differentially methylated genes (PANTHER Wnt signaling pathway, Bonferroni-corrected p < 2x10^-11^ and p < 3x10^-20^ for GSE58477 and GSE63409 respectively, see Supplementary Files 1 and 2). This suggests that, although we lose some resolution by combining CpGs by gene, we do capture major trends in the AML methylation landscape.

Having verified that gene-level consolidation provides similar results to CpG-level differential methylation, we applied ksRepo independently to each GEO dataset. 1,075 drugs had at least one overlapping gene with those annotated in 450K chips and were tested using ksRepo. We identified nine compounds with predicted FDR significance in only one of the datasets, and four that were significant in both. The four compounds significant in both datasets are presented in **Table 1** (results for all drugs are presented in Supplementary Table 1).

We posited that the most clear demonstration of utility for our method would be to correctly predict significance for the most commonly used AML therapy, cytarabine, which has remained the frontline therapy for over 30 years, and which is broadly effective against all subtypes of AML [31,32]. Indeed, our method did correctly identify cytarabine as highly significant in both datasets examined; however, our method did not identify either daunorubicin or idarubicin, two anthracycline chemotherapeutics that are commonly co-prescribed with cytarabine [31,33].

In addition to cytarabine, ksRepo nominated three novel compounds for use in AML: alitretinoin, panobinostat, and progesterone. Both alitretinoin, a geometric isomer of tretinoin currently used in the treatment of Acute Promyelocytic Leukemia (a subclass of AML) [34], and panobinostat, a histone deacetylase inhibitor commonly used in multiple myeloma [35], have been investigated for use in AML [36–39]. Lastly, while progesterone itself has not, to our knowledge, been suggested for AML treatment, a synthetic progestin, medroxyprogesterone acetate, has recently been suggested as a possible drug repositioning candidate for AML [40].

While our pipeline for epigenome-driven repositioning provided promising results for AML, it does have key limitations. First, as described above, removing the requirement for directionality (e.g. up- or down-regulation) provides substantial generality and allows for the use of CpG methylation data. However, such generalization does allow for the possibility that ksRepo candidates may have disease-exacerbating effects on their target genes; it is important, therefore, to perform *in vitro* and *in vivo* experiments to confirm which candidates are beneficial. Another possible drawback of non-directional repositioning is the possibility of missing promising candidates; however, as previous studies using directional repositioning techniques failed to capture significance for cytarabine, and predicted fewer plausible repositioning candidates than ksRepo [41–43]. A second limitation is the requirement that methylation or other epigenomic marks are annotated to a specific gene or genes; however, in the future, as our understanding of extragenic enhancers and other epigenomic features improves, this limitation should diminish. Further investigation into and annotation of other epigenetic modifications will also help to broaden the available inputs for our method. Lastly, we acknowledge that the CTD may contain a relatively small proportion of the true compound-gene interaction space, and future versions of ksRepo will incorporate other data sources to broaden the available databases included in the ksRepo package.

## CONCLUSIONS

In this study, we describe a pipeline for nominating drug repositioning candidates on the basis of differential methylation data. We combined a strategy for condensing CpG-level differential methylation to gene-level significance with our previously described drug repositioning tool, ksRepo. Unlike previously described methods of gene expression-based repositioning, which require up- or down-regulated gene or probe-level signals, our method is capable of leveraging complex gene-level differential methylation structure, for which there is often not a single directional effect. We applied our pipeline to discover repositioning candidates in AML, a disease with substantial epigenomic perturbations. We predicted significance for cytarabine, the most commonly prescribed AML medication, and identified three additional candidates with ongoing investigations for use in AML. Our method can be directly applied to other diseases with clear epigenetic perturbations and can be extended beyond DNA methylation to a variety of other epigenomic and molecular signals with complex or absent directionality.

## DECLARATIONS

### Ethics approval and consent to participate

Not applicable.

### Consent for publication

Not applicable.

### Availability of data and material

AML datasets are publicly available through the Gene Expression Omnibus (Accession #’s GSE58477 and GSE63409). The ksRepo R package is available from github <u>(https://github.com/adam-sam-brown/ksRepo)</u>.

### Competing interests

The authors have no competing interests to declare.

### Funding

ASB was supported by an National Institutes of Health (NIH) Training grant from the National Human Genome Research Institute (NHGRI), T32HG002295-12. CJP is supported by a National Institute of Environmental Health Sciences (NIEHS) R00 ES023504, R21 ES025052, a gift from Agilent Technologies, and a PhRMA fellowship. The funders had no role in study design, data collection and analysis, decision to publish, or preparation of the manuscript.

### Authors’ contributions

ASB and CJP conceived of the study. ASB conducted all statistical analyses. ASB and CJP wrote the manuscript.

## Acknowledgements

Not applicable.

